# Accurate simultaneous sequencing of genetic and epigenetic bases in DNA

**DOI:** 10.1101/2022.07.08.499285

**Authors:** Jens Füllgrabe, Walraj S Gosal, Páidí Creed, Sidong Liu, Casper K Lumby, David J Morley, Tobias W B Ost, Albert J Vilella, Shirong Yu, Helen Bignell, Philippa Burns, Tom Charlesworth, Beiyuan Fu, Howerd Fordham, Nick Harding, Olga Gandelman, Paula Golder, Christopher Hodson, Mengjie Li, Marjana Lila, Yang Liu, Joanne Mason, Jason Mellad, Jack Monahan, Oliver Nentwich, Alexandra Palmer, Michael Steward, Minna Taipale, Audrey Vandomme, Rita Santo San-Bento, Ankita Singhal, Julia Vivian, Natalia Wójtowicz, Nathan Williams, Nicolas J Walker, Nicola C H Wong, Gary Yalloway, Joanna D Holbrook, Shankar Balasubaramanian

**Affiliations:** Cambridge Epigenetix Ltd, The Trinity Building, Chesterford Research Park, Cambridge, UK; Cancer Research UK Cambridge Institute, University of Cambridge, Cambridge, UK; Yusuf Hamied Department of Chemistry, University of Cambridge, Cambridge, UK

## Abstract

DNA comprises molecular information stored via genetic bases (G, C, T, A) and also epigenetic bases, principally 5-methylcytosine (5mC) and 5-hydroxymethylcytosine (5hmC). Both genetic and epigenetic information are vital to our understanding of biology and disease states. Most DNA sequencing approaches address either genetics or epigenetics and thus capture incomplete information. Methods widely used to detect epigenetic DNA bases typically fail to capture common C-to-T mutations or distinguish 5mC from 5hmC. Here, we present a single-base-resolution sequencing methodology that will simultaneously sequence complete genetics and complete epigenetics in a single workflow. The approach is non-destructive to DNA and provides a digital readout of bases, which we exemplify by simultaneous sequencing of G, C, T, A, 5mC and 5hmC; 6-Letter sequencing. We demonstrate sequencing of human genomic DNA and also cell-free DNA taken from a blood sample of a cancer patient. The approach is accurate, requires low DNA input and has a simple workflow and analysis pipeline. We envisage it will be versatile across many applications in life sciences.

---

Information encoded in nucleic acids is fundamental to the biology of living systems. There are multiple dimensions of information stored within DNA. Genetic sequencing of the DNA bases G, C, T and A has been transformed by high throughput sequencing approaches in the past two decades. Epigenetic information in DNA provides insights into dynamic changes in biology that is closely associated with transcriptional programmes^1^ and cell fate^2^. The combination of genetic and epigenetic information provides a more comprehensive view of biology. The analysis of somatic genetic mutations, together with DNA methylation marks, from blood DNA gave a substantially more accurate prediction of mortality, than either genetics or DNA methylation alone could provide^3^. Both DNA methylation and genotype are required to determine the pluripotency of induced stem cells^4^ and their maturation capacity^5^. Germline genetic alterations cause changes in DNA methylation that ultimately dictates predisposition for disease^6,7,8^. Combining information on DNA methylation together with genetic sequence in cell-free DNA from blood, has been shown to substantially increase sensitivity to detect tumour DNA^9^. DNA methylation information can also inform on the tissue of origin of the tumour^10^. Non-invasive prenatal diagnostic analysis has also demonstrated that DNA methylation signal can determine foetal origin of DNA sequence^11^. Epigenetic information in DNA has been retrieved principally via sequencing 5-methylcytosine (5mC). More recently the 6^th^ base, 5-hydroxymethylcytosine (5hmC) has emerged as an important base modification that can provide information that goes beyond 5mC and genetics^12,13^. Hitherto researchers have accessed either genetic or epigenetic information, without resolving 5mC from 5hmC.

Commonly used sequencing approaches do not capture full information from both genetics and epigenetics. Next Generation Sequencing directly captures the canonical bases G, C, T and A in their readout^14^. A number of base conversion chemistries have been developed to help differentiate unmodified C from its epigenetic variants, 5mC or 5hmC. These include bisulphite-based approaches such as whole genome bisulphite-sequencing (WGBS)^15^ and also bisulphite-free approaches such as ACE-seq^16^, EM-seq^17^ and TAPS^18^. An important shortfall of all such methods is that conversion of either the C base, or one of its epigenetic derivatives, to a U (read as T) compromises the direct detection of genetic C-to-T changes, which is the most common mutation in the mammalian genome^19^ and in cancer^20^. Furthermore, the ambiguity caused by C-to-T conversions in the sequenced reads being mapped against either C or T in the reference genome increases false-positive matches and the search space, consequently making computational alignment and mapping of converted reads slower, more expensive and less accurate^21^. Furthermore, these existing methods cannot distinguish 5mC from 5hmC in a single workflow. Methods to distinguish 5hmC from 5mC by exclusively converting only one have been developed e.g. oxBS^22^, TAPSβ^23^, TAB-seq^24^ or by selectively copying 5mC across strands of DNA^25,26^. However, some of these can involve separate, parallel workflows and sequencing to yield full information, which may increase sample requirement, cost and time taken and/or yield data that lacks phased information. Combining separate data sets is fraught with difficulties that leads to additive measurement error and coverage gaps across workflows [see supplementary figure 1].

Third generation sequencing systems^27,28^ directly measure epigenetic and genetic modifications, in the same workflow. Such approaches necessitate machine learning to derive signal from noise in continuous measurement, and as such these analogue sequencing approaches have yet to achieve the context-independent base-level accuracy that a digital readout can produce. In DNA sequencing accuracy is critical and even the 1/1000 error rate achieved by Illumina sequencing is limiting for many applications^29^.

Herein we present a whole genome sequencing methodology capable of sequencing the 4 genetic letters in addition to 5mC and 5hmC to provide an accurate 6 letter digital readout in a single workflow. The processing of the DNA sample is entirely enzymatic and avoids the DNA degradation and genome coverage biases of bisulphite treatment^30,31^. The method uses all four genetic letters for genomic alignment and encompasses an intrinsic error suppression modality which leads to high accuracy for both genetic and epigenetic base calling. The approach is versatile and we demonstrate its application to different sample formats including human genomic DNA and a cell-free DNA sample from a human cancer patient.

We have developed a method to sequence beyond the four canonical DNA bases and include 5mC and 5hmC as the 5^th^ and 6^th^ DNA letters, respectively. The approach is compatible with any sequencer platform. Watson-Crick base pairing provides an explicit digital molecular mechanism for reading up to 4 letters (or states) [Fig 1b]. In order to read an epigenetic letter one can carry out a base conversion transformation, such as a C-to-T conversion (e.g. bisulphite seq/EM-Seq), where modified Cs are not converted [Fig 1b]. Here, the genetic information is compromised, and importantly masks the commonly occurring C-to-T single base variant. More generally, when sequencing DNA using a single-base coding system with a 4-state readout (i.e. G, C, T, A), one can at maximum unambiguously report on four states of information in a given run, be those genetic or epigenetic. A system that uses a two-base coding approach, whereby combinations of two bases relay the information for a state, enables up to sixteen states to be decoded unambiguously [Fig 1c]. This makes it possible to read all four genetic states and multiple epigenetic states in a single run. We have reduced this to practice for simultaneous 5-letter, and also 6-letter sequencing.

**Figure 1:**
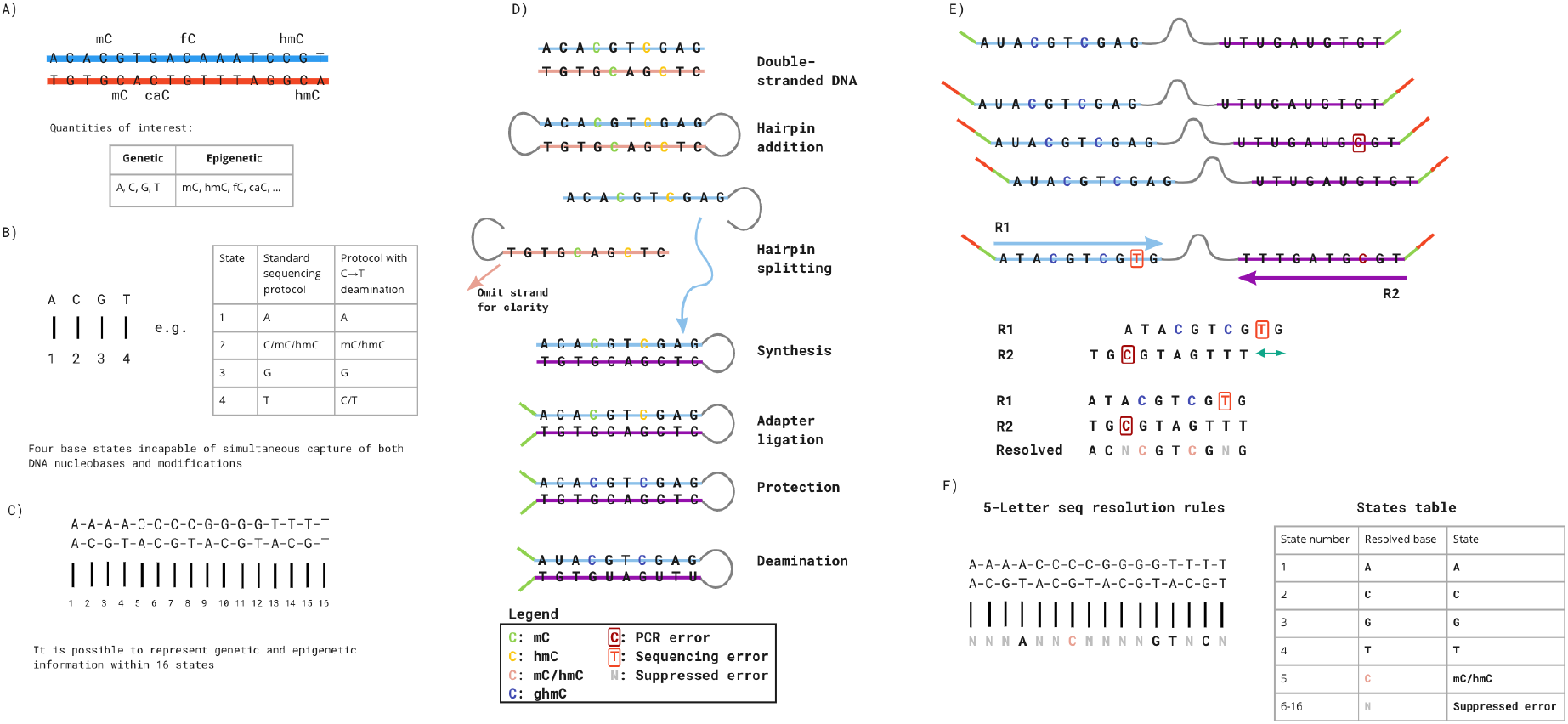
5-Letter seq **(A)** Double-stranded DNA with base modifications. **(B)** Traditional genetic sequencing only captures four states of information, which makes it impossible to determine genetic and epigenetic information. Base conversions can alter the information output, but the approach is inherently limited by only having four output states. **(C)** Two-base coding results in 4^2^ = 16 possible states enabling simultaneous determination of epigenetic and genetic states. **(D)** Laboratory workflow. Hairpins are ligated to double-stranded DNA and the strands are separated. The 3’-5’ strand is omitted for clarity, but follows a similar procedure to the 5’-3’ strand. The missing strand is synthesised and short adapters are ligated. Modified cytosines are protected. Unprotected Cs are converted via deamination, from C to U (read as T). **(E)** Sequencing protocol. The hairpin is opened and PCR amplified. Indexing adapters are added and the templates are paired-end sequenced. The two reads represent the same stretch of DNA and needs to be locally aligned to minimise errors. Using a set of resolution rules, the pairs of bases across the two reads are resolved into one of five states: A, C, modC, G, T. The method is able to identify errors occurring during PCR and sequencing. **(F)** Overview of the resolution rules and states under the 5-letter decoding model.

We first describe the 5-letter sequencing workflow that unambiguously resolves the four genetic bases with the fifth base being one of the epigenetic modifications, 5mC or 5hmC, termed hitherto as modified C or modC. In this workflow [Fig 1d], the sample DNA is first fragmented via sonication and then ligated to short, synthetic DNA hairpin adaptors at both ends. The construct is then split to separate the sense and antisense sample strands. For each original sample strand a complementary copy strand is synthesised by DNA polymerase extension of the 3’-end to generate a hairpin construct with the original sample DNA strand connected to its complementary strand, lacking epigenetic modifications, via a synthetic loop. Sequencing adapters are then ligated to the end. Modified cytosines are enzymatically protected. The unprotected Cs are then deaminated to uracil, which is subsequently read as thymine [Fig 1d]. The deaminated constructs are no longer fully complementary and have substantially reduced duplex stability, thus the hairpins can be readily opened and amplified by PCR. The constructs can be sequenced in paired-end format whereby read 1 (P1 primed) is the original stand and read 2 (P2 primed) is the copy stand. The read data is pairwise aligned so read 1 is aligned to its complementary read 2. Cognate residues from both reads are computationally resolved to produce a single genetic or epigenetic letter [Fig 1e]. Pairings of cognate bases that differ from the permissible five are the result of incomplete fidelity at some stage(s) comprising sample preparation, amplification, or erroneous base calling during sequencing. As these errors occur independently to cognate bases on each strand, substitutions result in a non-permissible pair. Non-permissible pairs are masked (marked as N) within the resolved read and the read itself is retained, leading to minimal information loss and high accuracy at read-level. The resolved read is aligned to the reference genome. Genetic variants and methylation counts are produced by read-counting at base-level.

The illustrative examples we provide below have deployed the sequence adaptors for the Illumina Novaseq NGS platform, however other adaptors can be readily substituted, and the method is compatible with any sequence reader capable of discriminating at least the 4 genetic bases.

We performed the 5-letter sequencing workflow on a mixed sample comprising 80 ng of sonicated human genomic DNA from a B-lymphoblast cell line (NA12878), obtained from the Genomes in a Bottle project, 0.4 ng bacteriophage λ DNA enzymatically methylated at all cytosines in CpG context by CpG Methylase and 0.4 ng pUC19 isolated from a methylation-negative strain of E.coli (Dam–, Dcm–). DNA was prepared, in duplicate, using the workflow outlined in Fig 1D (supplementary method 2) and sequenced on an Illumina Novaseq 6000 to produce approximately 550M paired-end reads (supplementary methods 3). These reads were computationally resolved as outlined in Figs 1 E, F (supplementary method 4). On average 97% of reads obtained were resolved and of these 90%_were aligned to the genome with mapq > 0. The average duplication rate in 5-Letter seq data was 10% with 90% of the genome covered by at least 1 read, and a 14x average coverage of the genome.

Resolved reads contain the 4 state genetic information (with the epigenetic information stored in .SAM format). Full genetic information enables genomic alignment using standard tools and reduced execution times compared to the 3-state alignment necessitated by techniques such as WGBS and EM-seq. The execution time for genomic alignment of 500,000 resolved 16-state 5-Letter seq reads using BWA-MEM was just under 8 minutes and the genomic alignment time for 500,000 3-state reads using BWA-Meth was 15.5 minutes.

We next compared the data quality of both the epigenetic and genetic components of 5-Letter seq, with best-practice-methods used to sequence either epigenetics only, or genetics only. To compare the epigenetic component of our method, the same sample mix (80 ng NA12878, 0.4 ng λ and puC19), was interrogated in duplicate by WGBS (Epitect, Qiagen) and by EM-seq (NEB), each with 275M paired-end reads. The data was processed in the standard methylseq pipeline, with 3-letter genomic alignment, yielding 89% and 92% aligned reads, and 15x and 17x deduplicated average coverage, for WGBS and EM-seq respectively. To compare the accuracy of the genetic sequencing component of our method, the same sample mix was interrogated by Illumina sequencing with standard library preparation using KAPA HyperPrep kit (Illumina) and processed in a similar pipeline (see supplemental methods). We calculated sensitivity to detect modC (expressed as a percentage) by considering all CpG-context Cs in the lambda genome and evaluating the ratio of modCs to the total number of observed cytosines (modified or unmodified). Similarly, specificity was calculated as the ratio of unmodified cytosines to the total number of cytosines (modified or unmodified) for CpG contexts in the pUC19 reference. Sensitivity of 5-Letter seq was 98.2% and specificity was 99.98%. This compared well to EM-seq which was less sensitive (97.89%) and less specific (99.5%), and to WGBS which was less sensitive (95.8%) and less specific (99.93%) [Fig 2a, top]. Across the genome average levels of modC observed at CHG and CHH sites were 0.09% as measured by 5-Letter seq, 0.13% as measured by WGBS and 0.32% as measured by EM-seq. In contrast, average modC levels at CpG sites was highest as measured by 5-Letter seq (53%), 52.17% as measured by EM-seq and 50.75% as measured by WGBS [Fig 2a, bottom]. Quantification of modC across reads and at genome level was highly concordant at single base level between 5-Letter seq and WGBS (Pearson’s p<0.01 r = 0.97) [Fig 2b]. There was a high level of agreement between modC estimates generated from the two methods [Fig 2c]. For the comparisons in Figures 2b and 2c, we pooled counts of modified and unmodified C calls at CpGs across both strands and across our two technical replicates, and then down-sampled to obtain 20x coverage for each technology.

**Figure 2:**
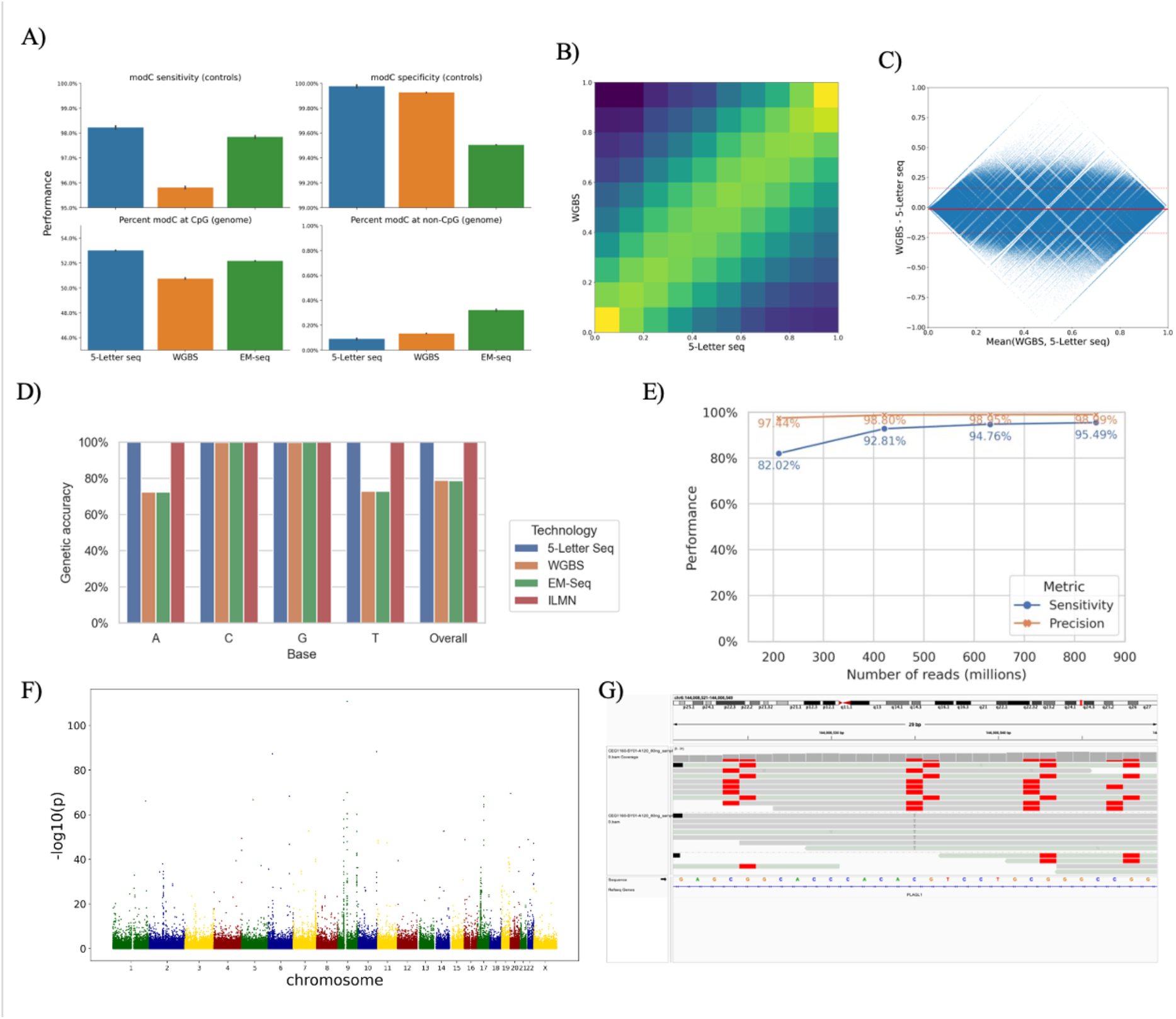
**(A, top)** Sensitivity and specificity of modC calling in 5-Letter seq, as computed on spike-in ground truth control sequences for 5-Letter seq (blue), WGBS (orange) and EM-seq (green); **(A, bottom)** average modC levels across all autosomes in NA12878 at CpG (left) and non-CpG (right) contexts; (**B)** Correlation heatmap showing high levels of agreement with WGBS (Pearson’s R 0.97);(**C)** Bland-Altman plot, with the average of the modC levels between the two methods on the x-axis and the difference on the y-axis (median difference of 1% with 95% of CpGs differing by between −16% and 22%, indicated by solid and dashed red lines, respectively); (**D)** Genetic accuracy as calculated on NA12878 high confidence regions for 5-Letter seq (blue), WGBS (orange), EM-seq (green) and standard Illumina sequencing (red); **(E)** Precision and sensitivity of variant calling using different quantities of 5-letter seq reads; (**F)** Manhattan plot of allele-specific methylation in NA12878. X-axis is chromosomal location and y-axis is −log10(p) from Fisher’s exact test of association between genotype and in cis modC levels; (**G)** Integrated genomic view (IGV) of 30nt region of PLAG1 gene in a C/T heterozygote. Reads to the C allele also show modC at the SNP site, and at CpGs −11nt, +7 and +12 nt from the SNP. Reads to the T allele show unmodified C at CpGs −11nt, +7 and +12 nt from the SNP

To evaluate the genetic component, called bases were compared to the reference at non-variant sites according to published variant call files for NA12878. Genetic accuracy was calculated as the ratio of correct base calls to total base calls (disregarding N calls). For 5-Letter seq and ILMN seq, the genetic accuracy was consistent across all base types (99.95%–99.98%). For WGBS and EM-seq, however, accuracy was high (99.69%–99.90%) for C/G bases, but low (72.42%-72.82%) for A/T bases [Fig 2d]. This is caused by unmodified C to T deamination in WGBS and EM-seq technologies that results in T bases being called instead of true C bases for reads mapping to the forward strand, and in A bases being called instead of true G bases for reads mapping to the reverse strand. This phenomenon has the consequence of masking actual C to T transitions. The accuracy of variant calling was investigated by applying GATK HaplotypeCaller to mapped read files of varying depths (211M-842M reads, resulting in mean coverages of 7X and 29X). Overall performance (SNPs and indels) were determined through comparison with published variant call files for NA12878. Sensitivity varied from 82% (211M reads) to 95% (842M reads) with precision consistently above 97% [Fig 2e].

It is important to note that 5-Letter seq simultaneously determines genetics and epigenetics on the same read. Therefore, combinations of genetic and epigenetic marks *in cis* on the same DNA molecule are detected. This property can be used to deconvolute genetic and epigenetic properties of heterologous samples. As an example, we detected allele-specific methylation in the NA12878 sample. Allele specific methylation is a wide-spread phenomenon in the human genome whereby DNA methylation is differential between alleles. It can identify regulatory sequence variants that underlie GWAS signals for common diseases^32,33^. Reads covering a polymorphism were interrogated for methylation levels that differed between alleles. Allele specific methylation was prevalent across the NA12878 genome [Figure 2f]. Loci with the strongest effects included *PLAG1*, a well-known imprinted gene^34^[Figure 2g].

The analysis of cell-free DNA (cfDNA) is a burgeoning aspect of diagnostics and application areas include enhanced non-invasive prenatal diagnosis, early cancer detection and disease monitoring^35^. A practical challenge is to work with the limiting amount of cfDNA that can be extracted from a standard blood draw, which is typically around 10 ng/ml^36^. An accurate sequencing method that can simultaneously detect genetic and epigenetic information from the same sample could transform cfDNA analysis. We deployed the 5-Letter seq workflow (Fig 1d-e) for the analysis of cfDNA from an individual with stage III colon cancer. Input DNA quantity was varied from 1 to 20 ng, in each case mixing the unsonicated sample with λ and pUC19 DNA spike-in controls, as previously described. Duplication rates ranged from 87% to 18% as input cfDNA increased from 1 ng to 20 ng respectively [Fig 3a], with genomic coverage remaining above 80% at all input levels and above or close to 90% for 5ng and up [Fig 3b]. The accuracy of methylation detection, determined for the λ and pUC19 DNA controls, remained consistently high at 98.5% sensitivity and 99.97% specificity (Fig 3c) even at 0.05 ng of control DNA in the 1 ng mixed sample.

**Figure 3:**
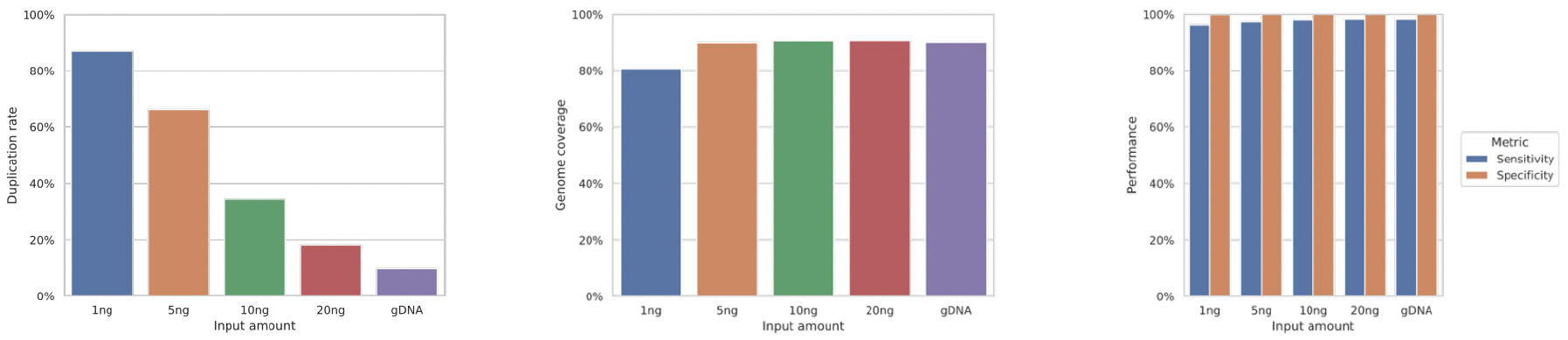
Application for 5-Letter seq to cfDNA. **(A)** Duplication rates achieved at variable input of cfDNA + 80ng gDNA **(B)** Proportion of genome covered with at least 1 read at variable inputs of cfDNA + 80ng gDNA **(C)** Sensitivity and specificity of modC detection is unaffected by input amount.

5-hydroxymethylcytosine (hmC) is the sixth DNA base and is generated by enzymatic oxidation of 5mC^37^. 5hmC has been shown to have value as a marker of biological states and disease which includes early cancer detection from cell-free DNA ^38,39^. We have adapted our platform to enable 6-Letter sequencing of DNA that comprises G, C, T, A, mC and hmC [Fig 4a]. A critical requirement is to disambiguate 5mC from 5hmC without compromising genetic base calling within the same sample fragment. The first three steps of the workflow are identical to 5-Letter sequencing shown in Fig 1d to generate the adapter ligated sample fragment with the synthetic copy strand. Methylation at 5mC is enzymatically copied across the CpG unit to the C on the copy strand, whilst 5hmC is enzymatically protected from such a copy. Thus, unmodified C, 5mC and 5hmC in each of the original CpG units are distinguished by unique 2-base combinations. The unmodified cytosines are then deaminated to uracil, which is subsequently read as thymine. The DNA is subjected to PCR amplification and sequencing as described earlier [Fig 1e]. The reads are pairwise aligned and resolved using the 2-base code shown in Fig 4b. Each of unmodified C, 5mC and 5hmC can be resolved as the three CpG units are distinct sequencing environments of the 2-base code [Fig 4b]. We estimated the accuracy of base modification status calls produced by 6-Letter seq using groundtruth control sequences with known modifications. We used the same fully unmethylated pUC19 and fully methylated lambda as were used in the evaluation of the 5-letter sequencing protocol and an additional short synthetic oligonucleotide with 5hmC present at specific CpGs. 6-Letter seq correctly identified 95.17% 5mC and 96.37% 5hmC status calls, at read level [Fig 4c] and retained a very high (99.98%) accuracy of detecting unmodified C at read level.

**Figure 4.**
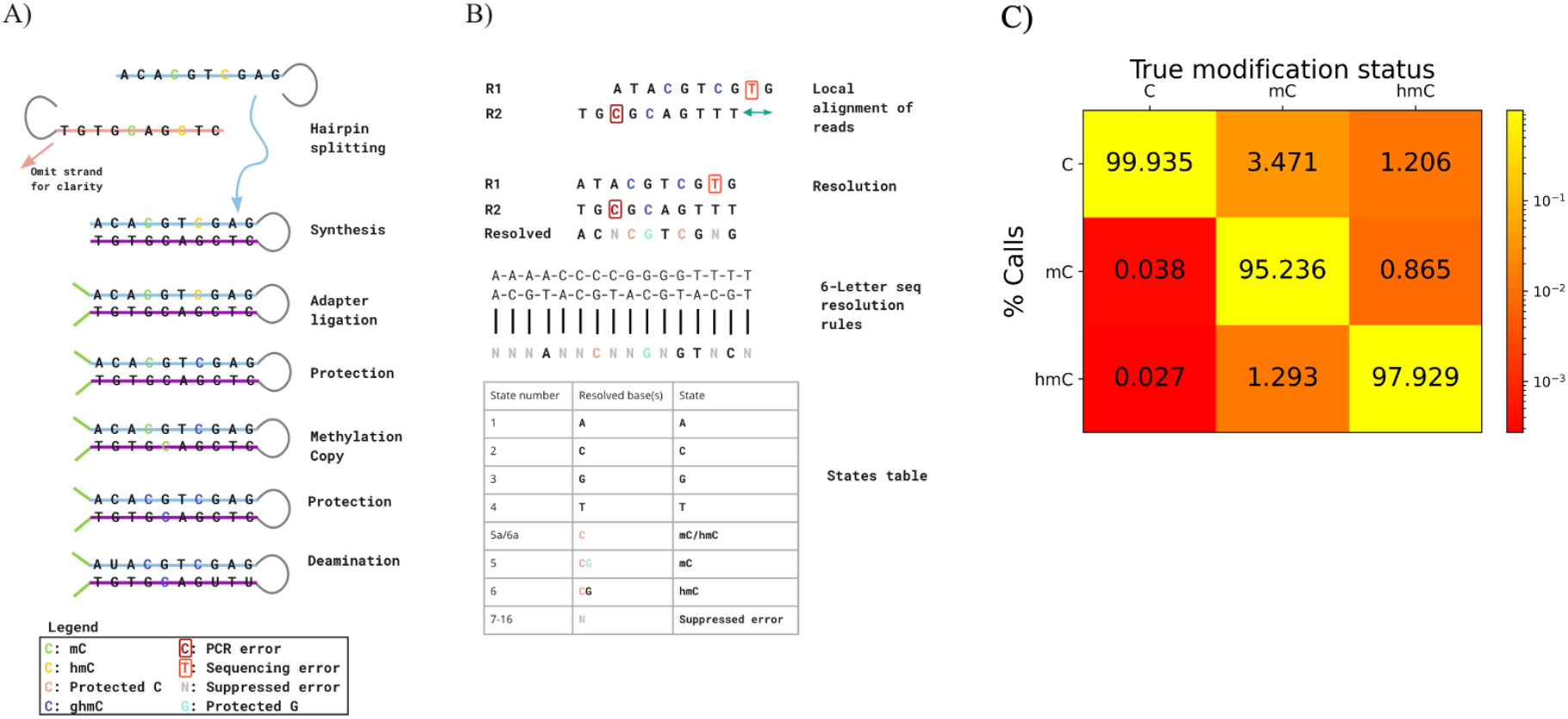
6-Letter seq **(A)** Schematics of 6-letter epigenetic sequencing protocol. A similar protocol to that of 5-Letter seq described in Fig 1D is followed with the addition of a methyl-copy step which copies the 5mC from the original to the copy strand. 5hmC is protected and not copied. **(B)** Overview of the resolution rules and states under the 6-letter decoding model. **(C)** Call-rate matrix, which contains the rate at which 6-Letter seq calls unmodified C/5mC/5hmC when the true state is unmodified C/5mC/5hmC. This is estimated from ground truth control sequences for which the modification status of each CpG is known. We calculate the rate at which 6-Letter seq calls unmodified C / 5mC / 5hmC at unmodified CpGs on a fully unmethylated pUC19 (first column), the rate at which 6-Letter seq calls unmodified C / 5mC / 5hmC at 5mCpGs on a fully methylated lambda (second column), and the rate at which 6-Letter seq calls unmodified C / 5mC / 5hmC at 5hmCpGs on a synthetic oligonucleotide (third column).

We have described a sequencing platform that will deliver the full complement of genetics in addition to the epigenetic bases 5mC and 5hmC at base resolution in a single workflow and data pipeline. The platform operates via a two-base coding mechanism at the molecular level coupled with decoding software, which also improves the accuracy of genetic sequencing and variant calling alongside high-accuracy of epigenetic base calling. The platform comprises an all-enzyme workflow with extremely high conversion efficiencies thus enabling accurate data from valuable, low input biological samples or from clinical cell-free DNA. The readout of genetic or epigenetic bases is digital and context-independent, without requirement for learning or training algorithms. The alignment uses a 4-base system which is substantially faster and more accurate than 3-base alignments used for bisulphite sequencing or EM-Seq. The acquisition of accurate, phased genetic and epigenetic information, in a single experimental and data workflow is likely to offer numerous advantages in research and in diagnostics by providing more comprehensive biological information, with ease of use and at reduced cost. As the platform fundamentally utilises Watson-Crick base pairing to decode information, it can be made readily compatible with any sequencer platform, besides Next Generation Sequencing and we see opportunities for its future application to long-read sequencing and single cell analysis.

## Methods

### Samples

Genomic DNA was obtained from the Coriell Institute (https://www.coriell.org/0/Sections/Search/Sample_Detail.aspx?Ref=NA12878)

Double spun plasma was obtained from Trans-hit Biomarkers Inc (Laval, Quebec, Canada) from a single male patient with a CRC diagnosis. cfDNA was extracted using the NextPrep-Mag™ kit on the Chemagic Prime platform (Perkin Elmer chemagen Technologie GmbH, Baesweiler, Germany). Fully methylated lambda and unmethylated pUC19 controls were generated as per supplementary method 1

### Laboratory processing

80ng of sonicated genomic DNA and 10ng cfDNA was processed by either 5-Letter Seq (Cambridge Epigenetix), EM-Seq (Cat#E7120, New England Biolabs) or EpiTect Plus DNA Bisulphite kit (Cat#59124, Qiagen) 5-Letter seq and 6-Letter seq was performed with kits available from Cambridge Epigenetix. EM-seq was performed with a kit available from New England Biolabs (Cat #E7120, NEB). WGBS was performed with the EpiTect bisulphite sequencing kit available from Qiagen (Cat #5912. ILMN sequencing was carried out with the KAPA Hyper Prep kit (Cat #KK8500, Roche). Deviations from the manufacturer’s instructions are described in supplementary methods 2. Samples were sequenced on Illumina Novaseq. (supplementary methods 3 for sequencing conditions)

### Informatics

FASTQ files were downloaded from BaseSpace and processed using software created by Cambridge Epigenetix [supplementary methods 4]. Variant calling statistics were produced as described in supplementary methods 5. Production of the call rate matrix for 6-letter sequencing is described in supplementary methods 6.

