## Supplementary Materials for "Accurate simultaneous sequencing of genetic and epigenetic bases in DNA"

### Supplementary Methods 1

Ground truth controls

An 80 bp oligonucleotide with balanced CG content and no homology to human genome, lambda or pUC19, and symmetrical 5hmC at one CpG and asymmetrical 5hmC at another, was designed and purchased from ATDBio Ltd. The oligonucleotide pair was diluted in 100 mM Potassium Acetate; 30 mM HEPES, pH 7.5 (IDT) and annealed by heating to 94°C for 2 minutes and cooling gradually to 4°C.

```
5’pGTATTGACAAGGTGACAGTATCGTCCAGGGACAGTCTCGTAGTACCACCTAGTCTACTCGAGAATGTCAAGGTGTCAGAC  3’
3’ CATAACTGTTCCACTGTCATAGCAGGTCCCTGTCAGAGCATCATGGTGGATCAGATGAGCTCTTACAGTTCCACAGTCTGp 5’
```

C=5hmC
C=CpG

For unmethylated pUC19 control, the plasmid (NEB) was transformed into *dcm-/dam- E. coli* chemically competent cells (NEB). Plasmid was isolated from culture using a QIAprep Spin Miniprep Kit (Qiagen). Methylated lambda DNA was prepared using unmethylated lambda DNA (Promega). Briefly, 1 µg DNA was incubated with 8 units of CpG methyltransferase MSssI (NEB), NEB Buffer 2, 320 µM S-adenosyl-methionine (NEB) at 37°C. After 4 hours, a further 2 units of MSssI was added and the reaction incubated for a further 4 hours at 37°C. The DNA was purified using a Zymo clean and concentrator column (Zymo). Complete methylation was checked using methylation sensitive HpaII (NEB) and methylation insensitive MspI (NEB). A negative control using unmethylated lambda DNA but no MSssI was also prepared and used to check digestion with MspI and HpaII. Both methylated lambda DNA and unmethylated pUC19 were diluted in 10mM Tris-HCl pH 8.0 0.1mM EDTA and fragmented to ~250 bp using a Covaris M220.

**Supplementary Methods 2**

For 5-Letter Seq and 6-Letter Seq variable amounts of input material were used according to the manufacturers protocol (Cambridge Epigenetix). For input DNA either pre-sonicated genomic DNA (Coriell, Cat#NA12878) or cfDNA from a single colorectal cancer patient was used that was extracted from double spun plasma purchased from Trans-hit Biomarkers Inc (Laval, Quebec, Canada). Genomic DNA was sonicated using the Covaris M220 Sonicator set to a target size of 250bp.

To generate standard genomic libraries the KAPA Hyper Prep kit (Cat #KK8500, Roche) was used according to the manufacturer protocol with the following modifications. 100ng of genomic DNA (Coriell, Cat#NA12878) was sonicated to ~250 bp using a Covaris M220 sonicator in 50µl sonication tubes (Covaris) and used in the experiment. For adapter ligation, standard TRUSeq adapters were used (15µM, Illumina). Ampure beads (Beckman Coulter) were used instead of KAPA clean-up beads and the elution volume was reduced to 21µl using library dilution buffer (10mM Tris-HCl, pH8.0). For library amplification the KAPA Hifi Hot start PCR kit was used (Cat#KK2500, Roche) according to the manufacturers protocol using 4 PCR cycles and 1µM final primer concentration. Final libraries were quantified using D1000 HS screen tapes (Agilent).

EM-seq libraries were produced using the New England Biolabs EM-Seq kit (Cat# E7120, NEB) according to the manufacturers protocol with the following modifications. Instead of using the EM-Seq Control DNA, the ground truth spike-in DNAs described in Supplementary methods 1 were used to allow a direct comparison Denaturation was performed using Formamide (Cat# F9037, Sigma). For the final PCR, 7 cycles of PCR were used (80ng gDNA input). Libraries were quantified using D1000 HS screen tapes (Agilent) on a Tapestation (Cat# 4200, Agilent).

To prepare libraries for WGBS the libraries were prepared using the EM-Seq kit (Cat# E7120, NEB) and the EpiTect Plus DNA Bisulphite kit (Cat# 59124, Qiagen) with the following modifications. Instead of using the EM-Seq Control DNA, the ground-truth spike-in DNAs described in Supplementary methods 1 were used to allow a direct comparison. The EM-Seq protocol was followed until Clean-Up of Adaptor Ligated DNA and the eluate was then used as input for the EpiTect Plus DNA Bisulphite kit. The high-concentration sample setup was used for Bisulphite conversion. To improve Bisulphite conversion, the 20µl elution from Bisulphite conversion was used for another round of conversion using the EpiTect Plus DNA Bisulphite kit (Cat# 59124) again. After the second round of conversion the DNA was PCR amplified using the EM-Seq kit according to manufacturer's protocol starting from PCR Amplification using 8 cycles of PCR amplification.

**Supplementary Methods 3**

Sequencing was performed using the S4 Standard workflow on the Illumina NovaSeq. Libraries were quantified using the Tapestation D5000 (Agilent) and subsequently diluted to 1.2-1.5nM for loading on the sequencer. To balance out the low Cytosine content in deaminated libraries (WGBS, EM-Seq, 5-Letter Seq and 6-Letter Seq), 10% PhiX (Cat# FC-110-3001, Illumina) was added according to manufacturer protocol. All libraries were run in a Paired End set up using 200 cycle kits in a 111/8/8/111 base reads setup.

**Supplementary Methods 4**

Processing of 5-Letter seq and 6-Letter seq data

FASTQ files were trimmed and quality-filtered using fastp[40] before being processed through a resolution algorithm designed by Cambridge Epigenetix. Resolved FASTQ files were aligned using bwa mem[41] to a standard 4-letter reference genome comprising of both GRCh38 and spiked-in control sequences. Epigenetic information encoded in tags in the resolved FASTQ files was passed on into the aligned BAM files. The aligned BAM files were then split into reads aligning to the genome and reads aligning to the controls; unmapped reads were filtered out. Reads aligning to the genome, to the methylated bacteriophage lambda control, and to the unmethylated pUC19 control were deduplicated using Picard MarkDuplicates[42]. A range of standard metrics were calculated on the genome-aligned reads using samtools[43], Qualimap[44], deepTools[45] and Picard[46]. Accuracy of the genetic base calling was calculated relative to the known genotype of high-confidence regions of chromosome 20 of the NA12878 sample.

Quantification of epigenetic modifications was calculated at each CpG, CHG, and CHH site that was present in the reference genome and covered in the sequencing. This was performed using software developed by Cambridge Epigenetix. Likewise, quantification of epigenetic modifications was calculated at each CpG site in the methylated bacteriophage lambda control and the unmethylated pUC19 control. Sensitivity of modification calling was calculated from the methylated bacteriophage lambda control and specificity of modification calling was calculated from the unmethylated pUC19 control.

All of the processing was performed using a software pipeline developed by Cambridge Epigenetix, written in the Nextflow orchestration language and processed on the Google Cloud Platform (GCP).

**Processing of EM-Seq and WGBS Samples**

Processing of EM-Seq and WGBS samples was performed using a software pipeline written in the Nextflow orchestration language and processed on the Google Cloud Platform (GCP). Trimming of FASTQ files was performed using Trim Galore![46]. Alignment to a deaminated reference genome comprising of both GRCh38 and spiked-in control sequences was performed using bwa meth[47] and deduplication was performed using Picard MarkDuplicates. Modification-calling at CpG, CHG, and CHH sites was performed using MethylDackel[48].

**Processing of ILMN Samples**

Processing of Illumina samples was performed using a software pipeline written in the Nextflow orchestration language and processed on the Google Cloud Platform (GCP). Trimming was performed using fastp. Alignment to a GRCh38 reference genome was performed using bwa mem and deduplication was performed using Picard MarkDuplicates.

**Supplementary Methods 5 (variant calling)**

To examine variant calling performance, the 5-Letter seq reads from two 80ng gDNA (NA12878) samples were pooled using `samtools merge` (v. 1.15.1). The combined .bam file had a total read count of 842 million reads, equivalent to a mean coverage of 28.6X. The combined .bam file was then downsampled into four separate .bam files representing fractions of 0.25, 0.50, 0.75 and 1.0 times the original depth. This was achieved using `samtools view --subsample {fraction} --subsample-seed 1` and resulted in .bam files of 211M reads (7.1X), 421M reads (14.3X), 632M reads (21.5X), and 842M reads (28.6X) respectively. Variant calling was then performed for each of these samples by GATK HaplotypeCaller (v. 4.2.5.0) with default settings. For efficiency, HaplotypeCaller was run in parallel for each chromosome and the resulting VCF files were merged using GATK MergeVCFs. Finally, RTG Tools' vcfeval function (v. 3.12.1) was used to compare the 5-Letter seq variant call data to a ground truth set defined by the Genome in a Bottle VCF file for NA12878 (https://ftp-trace.ncbi.nlm.nih.gov/giab/ftp/release/NA12878_HG001/NISTv3.3.2/GRCh38/). Evaluation was constrained to the high confidence regions defined by the associated NA12878 .bed file (same location). Overall variant calling performance, representing performance across both SNPs and indels, was extracted from the summary.txt output file. Reported sensitivity and precision metrics were associated with ROC score thresholds resulting in maximal F-measure.

**Supplementary Methods 6** – call-rate matrix

A *call-rate matrix* is used to measure the accuracy of modified Cytosine calls. Each cell represents the rate at which the method calls a particular modification X when the true modification is Y, e.g., the cell M[C, mC] represents the rate at which a method calls *unmodified C* when the true modification status is *methylated C*. Each column of the matrix, corresponding to the rate at which we call each modification status for a particular true modification status, is estimated using a different spike-in control: a fully unmethylated pUC19 for the column corresponding to a true state of unmodified C; a fully methylated lambda for the column corresponding to a true state of methylated C; and a synthetic oligonucleotide for the column corresponding to a true state of hydroxymethylated C. For a given column, the rate is calculated at the proportion of bases with each modification status in the set of bases for which the genetic base call is C and which are aligned to CpGs in the given control sequence.

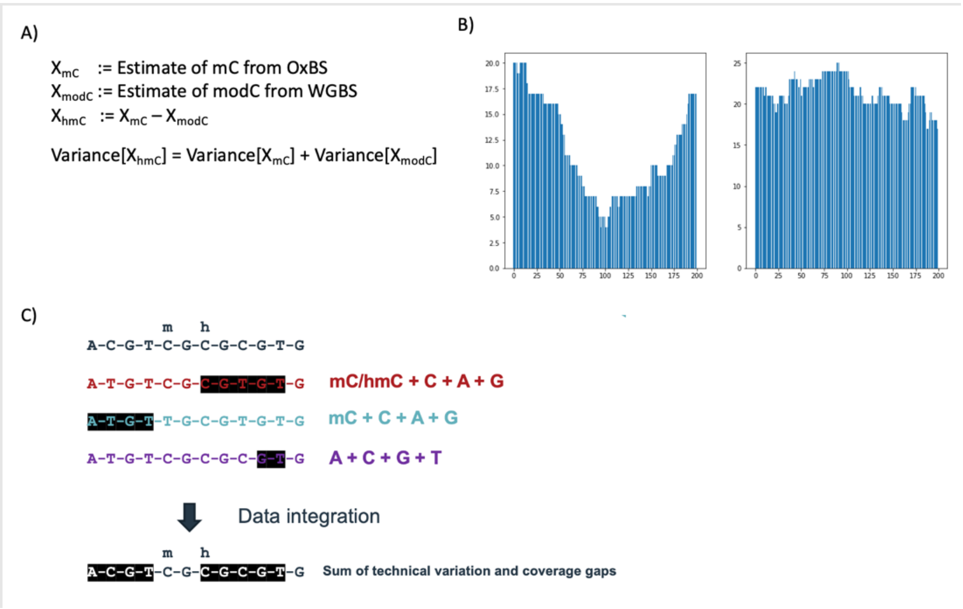


**Supplementary Figure 1** To obtain genetics, mC and hmC using existing methods, it is necessary to combine information from a standard whole genome sequencing experiment with data from separate WGBS and OxBS experiments. This presents a number of practical challenges. **SFig 1A)** illustrates how the variance in estimates for hmC levels at a given CpG is the sum of the variance of our estimates of modC from WGBS and mC from OxBS. **SFig1B)** contains an example of expected coverage (reads sampled uniformly from a genome) at the same 200nt region across two sequencing experiments with average 20x coverage; the left is a WGBS experiment and the right is an oxBS experiment. Whenever the coverage dips below 10x for either of the two – no call on mC or hmC can be made. **SFig1C)** provides a schematic illustrating how areas of low coverage can combine unfavourably when combining data across multiple experiments. The integrated dataset will have a coverage gap in any region which has insufficient coverage in any one of the experiments being combined.

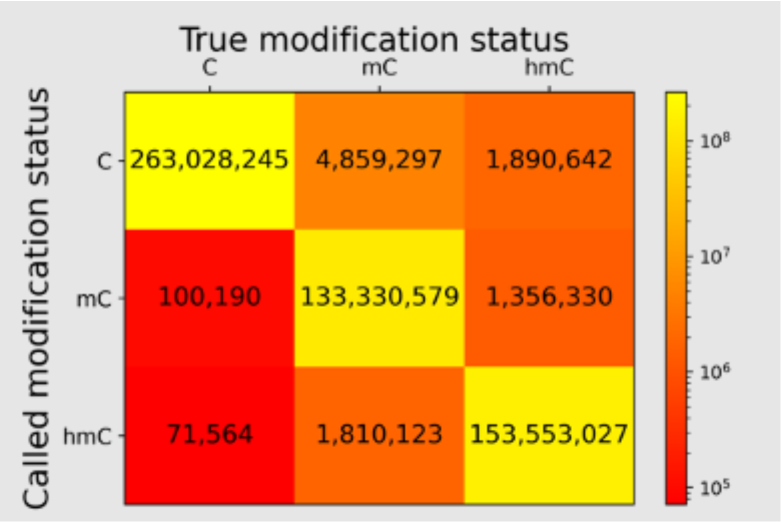


**Supplementary Figure 2** Given the estimated call-rate matrix for 6-letter sequencing, we can extrapolate to a genome with 56 million CpGs with relative abundances of a 5hmC rich genome like that of neurons[49] where 5hmC is believed to occur at approximately 28%, 5mC at approximately 25%, and unmodified C at approximately 47%. In this setting, we would expect to obtain highly accurate base modification calls with 98.8% of hmC calls occurring at sites which are indeed hydroxymethylated, 98.9% of mC calls occurring at sites which are indeed methylated, and 97.5% of unmodified C calls occurring at sites which are indeed unmodified.
